# Oscillatory and aperiodic neural dynamics shape temporal perception and weighting of perceptual priors in ADHD profiles

**DOI:** 10.64898/2026.01.30.702812

**Authors:** Gianluca Marsicano, Michele Deodato, David Melcher

## Abstract

Attention-deficit/hyperactivity disorder (ADHD) is characterized by an atypically compressed sense of time, yet the neural mechanisms underlying this atypical temporal perception remain poorly understood. Temporal perception depends on the brain’s ability to organize sensory input into coherent experiences, ensuring perceptual stability despite uncertainty. Oscillations in the alpha band (8–13 Hz) and aperiodic 1/f dynamics have been proposed as key neural mechanisms through which the visual system orchestrates sensory information over time. Individuals with ADHD show atypicalities in these neural dynamics, but how these features relate to ADHD differences in temporal processing remains unexplored. Here, we combined a sustained visual temporal integration task with resting-state EEG to test whether oscillatory and aperiodic neural dynamics jointly account for temporal processing performance across neurotypical participants with self-reported ADHD traits (n = 83). Higher ADHD features in both inattentive and hyperactive-impulsive domains were associated with narrower temporal binding windows and reduced serial dependence on prior perception, indicating sharper temporal resolution but diminished perceptual stability. Resting-state EEG revealed systematically faster individual alpha frequency (IAF) and flatter aperiodic spectra in individuals with higher ADHD traits. Mediation analyses showed that faster IAF explained ADHD-related reductions in temporal integration thresholds and serial dependence, whereas increased neural noise selectively amplified perceptual history-dependent biases. These findings reveal distinct oscillatory and aperiodic neural pathways through which ADHD features shape temporal perception, suggesting a multidimensional neural architecture underlying atypical temporal processing in ADHD.

## Introduction

Temporal perception constitutes a core function of the nervous system, supporting the integration of dynamic sensory information over time into coherent representations of the external world. Atypical temporal perception is widely observed in psychiatric and neurodevelopmental disorders, with disruptions in temporal processing implicated in higher-order cognitive deficits and atypical perceptual experiences^1,2,3,4,5,6,7,8,9,10,11^. An emblematic case is attention-deficit/hyperactivity disorder (ADHD), a neurodevelopmental condition characterized by inattention, hyperactivity, and impulsivity, in which atypical temporal perception has been proposed as a central component of its neurocognitive architecture^12,13,14,15,16^. Growing evidence converges to indicate that individuals with ADHD experience a contracted sense of time, perceiving the passage of time as unusually fast^15^. This alteration in temporal experience is thought to contribute to the peculiar profile of this disorder, including difficulties with sustained attention, heightened arousal levels, and impulsive behaviour^14,15^.

Converging findings on altered activity in sensory cortical areas strongly suggest the presence of a pure temporal deficit in ADHD, likely arising from low-level sensory processing anomalies^17,18,19,20,21^. Consistent with this view, recent evidence shows that individuals with ADHD exhibit a narrower window of temporal integration across sensory domains, tending to overestimate temporal asynchrony and judging significantly fewer stimuli as simultaneous^18,19,22,23,24^. Such perceived temporal misalignment across sensory events has been proposed to contribute to the characteristic behavioural phenotype of distractibility and inattentiveness observed in this condition^22,23^, as well as in neurotypical individuals displaying higher levels of ADHD-like traits^23,24^.

An influential empirical framework suggests that rhythmic neural activity in the alpha band (8–13 Hz) provides a temporal scaffold for perception^25^. According to this view, alpha frequency serves as a sampling rate for the sensory system: faster individual alpha rhythms support finer temporal resolution, leading stimuli that occur within the same alpha cycle to be bound into a single perceptual event^25^. In line with this view, recent evidence shows that faster alpha activity is associated with longer perceived durations^26^ (Mioni et al., 2020), narrower temporal integration windows^27,28,29^, and faster sampling rate of sensory information^29,30,31,32,33^. Despite the documented alterations, the relationship between alpha oscillatory activity and temporal perception in ADHD remains unexplored. Given the contracted sense of time and the self-reported feeling of being “*out of sync*” ^34^, it is essential to determine whether and how alpha oscillatory dynamics contribute to temporal processing atypicalities in ADHD.

In addition to oscillatory neural activity, temporal perception depends critically on non-rhythmic neural dynamics^27,28,29,35^. Indeed, aperiodic *1/f* neural activity^27,28,29,36,37^ can shape how oscillations organize sensory information over time. Flatter neural spectra, indicating greater neural noise, are hypothesized to disrupt alpha-mediated pulsed inhibition, weakening temporal coupling and leading to more variable and less precise temporal judgments^27,28,29,36,37,38^. This is particularly relevant for ADHD, as recent work shows that individuals with ADHD exhibit a markedly lower aperiodic exponent^39,40^, potentially linked to alterations in dopaminergic and noradrenergic systems often reported in this condition^41,42,43,44^. Accordingly, investigating how oscillatory and aperiodic neural dynamics jointly shape temporal processing in ADHD may provide a more complete account of the neural mechanisms underlying temporal atypicalities in this disorder.

Critically, the fundamental mechanisms linking temporal processing to the ADHD phenotype remain poorly understood. From a Bayesian perspective, temporal perception reflects the balance between sensitivity to incoming sensory evidence and the influence of internal models, with serial dependence on recent perceptual experience playing a central role in biasing the interpretation of new input^45,46^. Consistent with this view, events that follow a temporal binding percept are more likely to be experienced as a single unified event^7,29,47,48^. Recent work suggests that core ADHD symptoms, including impulsive-hyperactive response patterns and difficulties in sustaining attention, may reflect excessive precision assigned to incoming sensory information^48,49,50,51,52,53,54,55,56,57,58,59,60^. Yet, no systematic investigation has examined the influence of recent perceptual history over short timescales in ADHD profiles, its relationship to periodic and aperiodic neural activity, or how these processes influence their temporal perception.

At the neural level, emerging evidence indicates that both the speed of alpha oscillations and aperiodic components determine how strongly past perceptual experience shapes current temporal judgments: slower alpha rhythms and increased neural noise reduce sensory precision, potentially increasing reliance on prior experience^28,29,30^. Given the heightened attentional fluctuations and faster decay of information over short time scales reported in ADHD, it is conceivable that reduced integration of recent perceptual information into current decisions may arise from atypical alpha oscillatory and aperiodic neural dynamics.

Here, we investigated how oscillatory and aperiodic neural dynamics support multiple aspects of temporal processing in a large sample of individuals (n = 83) exhibiting varying degrees of ADHD features within the general population. To this end, we implemented a sustained visual integration paradigm^29^ (Fig. 1A) designed to estimate individual temporal integration thresholds over short time scales. Critically, the experimental design avoids potential confounds in temporal integration tasks related to attentional fluctuations that are characteristic of ADHD. This design allowed us to estimate individual temporal integration thresholds while avoiding confounds associated with traditional paradigms, such as detection failures, thereby providing a more direct measure of perceptual integration^29,61^. One advantage of continuously presented stimulus streams is that integration performance is not confounded by failures to detect rapidly presented stimuli. We recorded resting-state EEG in both eyes-open and eyes-closed conditions to examine how oscillatory and aperiodic neural dynamics jointly shape visual temporal integration in relation to ADHD profiles. We also assessed how perceptual decisions from the previous trial influence current temporal judgments, linking short-term perceptual history to periodic and aperiodic neural activity. Together, these measures revealed a mechanistic association in which higher ADHD features were predicted by faster alpha oscillatory speed and higher-noise neural background, accounting for both a narrower temporal binding window and a reduced influence of prior perceptual information.

**Figure 1.**
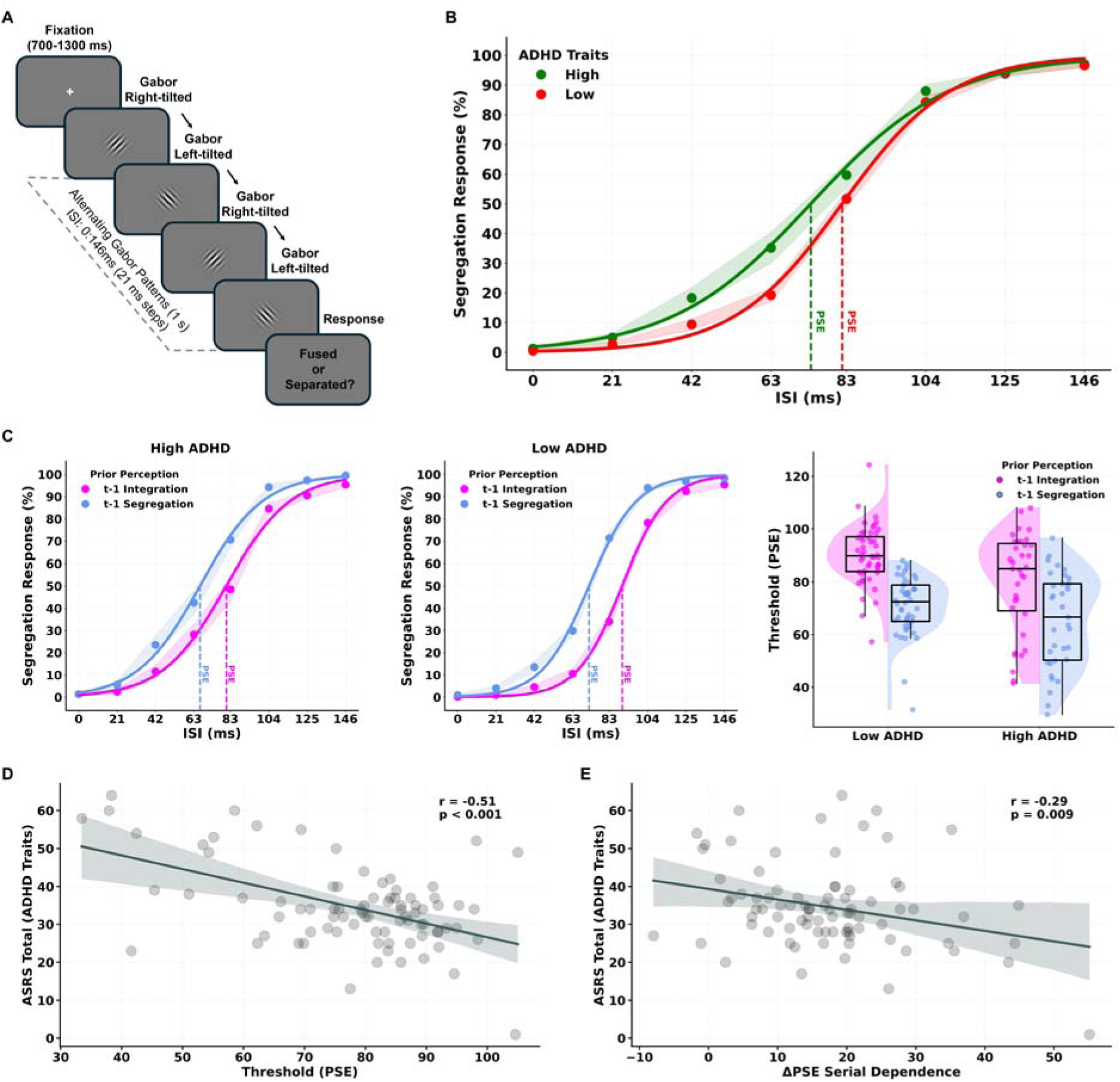
Atypical visual temporal processing in ADHD-like profiles. (A) Schematic depiction of the sustained-stream temporal integration task^29^. On each trial, two low–spatial-frequency Gabor patches of opposite orientation (±45°) alternated in counterphase for ∼1 s with variable inter-stimulus intervals (ISIs, 0–146 ms, steps of 21 ms). Participants reported whether they perceived a fused plaid (integration) or separate gratings (segregation), followed by a confidence rating (1–4). (B) Psychometric functions depicting segregation responses as a function of ISI for individuals with low (red) and high (green) ADHD traits (median split). Higher ADHD traits were associated with a leftward shift of the psychometric curve, reflecting lower temporal integration thresholds (PSEs) and therefore a narrower temporal binding window. Shaded areas denote SEM. (C) Individual PSE values as a function of previous perceptual outcome. Left and middle panels: psychometric functions separately for trials preceded by integration (t–1 integration; purple) or segregation (t–1 segregation; blue) responses in high-trait (left) and low-trait (middle) ADHD groups. Shaded areas denote SEM. Right panel: distribution of PSEs for each history condition, illustrating reduced serial dependence (smaller ΔPSE) in individuals with higher ADHD traits. Black boxplots indicate the median and interquartile range (IQR), and individual dots represent participant-level data. (D) Across participants, higher ADHD traits (ASRS total scores) were associated with lower temporal integration thresholds (PSE), indicating sharper visual temporal resolution. Shaded bands represent 95% confidence intervals. (E) Higher ADHD traits were also associated with reduced serial dependence (ΔPSE), reflecting a weaker influence of prior perceptual outcomes on current temporal judgments. Shaded bands represent 95% confidence intervals. Median-split visualizations in panels B-C are presented for illustrative purposes only; all statistical analyses treated ADHD traits as continuous variables.

## Results

### Sharper visual temporal perception and weaker perceptual bias in ADHD profiles

As described in our previous study^29^, the sustained stream of visual patterns in our temporal integration task elicited the illusory perception of a single fused plaid at shorter inter-stimulus intervals (ISIs), whereas longer ISIs induced the perception of two alternating gratings (segregation; Fig. 1A). To examine how variability in ADHD profiles, measured with the ASRS questionnaire^62^ (see Methods), relates to temporal integration performance, we computed for each participant the proportion of segregation responses across ISIs and fitted a logistic psychometric function. This yielded two key parameters: the 50% threshold (Point of Subjective Equality, PSE) and the slope. We correlated ASRS scores with PSE and slope values for the total ASRS score, the inattentiveness subscale, and the hyperactivity–impulsivity subscale. In line with evidence of faster temporal perception in ADHD^15^, we found that a narrower temporal binding window, reflected by lower PSE values, was associated with higher ADHD traits in total ASRS scores (r = −0.51, 95% CI [−0.65, −0.33], p < .001; Fig. 1D), as well as in the inattentiveness (r = −0.435, 95% CI [−0.59, −0.46], p < .001) and hyperactivity–impulsivity (r = −0.503, 95% CI [−0.64, −0.55], p < .001) subscales. No significant associations emerged between psychometric slope values and ADHD traits (all p > .389). Building on recent evidence that visual temporal processing is dynamically modulated by recent perceptual experience^7,29,61^, we next investigated whether ADHD traits influence the extent to which previous perceptual judgments bias current temporal perception. To quantify this serial dependence effect, trials were sorted according to the percept experienced on the previous trial (t–1 integration or t–1 segregation) and separate psychometric functions were fit for each condition. For each participant, the magnitude of serial dependence was defined as the difference between the two PSE estimates (ΔPSE = PSE t–1 integration – PSE t–1 segregation). Correlational analyses revealed that individuals with higher ADHD traits showed a weaker influence of prior perceptual experience on current temporal judgments, as reflected in reduced ΔPSE values for higher ASRS total scores (r = −0.286, 95% CI [−0.47, −0.097], p = .009; Fig. 1E), as well as for inattentiveness (r = −0.229, 95% CI [−0.42, −0.01], p = .038) and hyperactivity–impulsivity (r = −0.298, 95% CI [−0.48, −0.08], p = .006) subscales.

Taken together, these findings show that individuals with higher ADHD features exhibit atypical temporal perception, characterized by a narrower temporal binding window and reduced reliance on prior perceptual outcomes. This pattern suggests reduced flexibility in adaptively shaping temporal processing based on recent experience and reflects an altered dynamic interplay between sensory evidence and perceptual priors in individuals with elevated ADHD traits. All analyses treated ADHD measures as continuous variables, preserving variance and statistical power while avoiding the information loss associated with dichotomization^63^. Median-split groups shown in Fig. 1B–C were used solely for visualization and not for statistical inference (see Methods).

### Oscillatory and aperiodic neural dynamics in ADHD profiles

Alongside the temporal integration task, we recorded resting-state EEG activity (eyes-closed, EC; eyes-open, EO) from 64 scalp electrodes. To examine how alpha oscillatory speed and aperiodic neural dynamics vary with ADHD traits, we estimated each participant’s oscillatory spectrum and aperiodic components^64^ (see Methods and Supplemental Information S1).

We first correlated resting-state individual alpha frequency (IAF) with ASRS total scores and the inattentiveness and hyperactivity–impulsivity subscales, using Pearson correlations with cluster-based permutation correction (10,000 permutations) across all electrodes for both EC and EO conditions. Higher ADHD traits were significantly associated with faster IAFs, particularly over posterior–central scalp regions. Significant positive clusters emerged for IAF and ASRS total scores (EO: p < .001; EC: p = .018; Fig. 2A, left), hyperactivity–impulsivity (EO: p < .001; EC: p < .001; Fig. 2A, middle), and inattentiveness (EO: p = .026; EC: p = .005; Fig. 2A, right). We next examined how ADHD traits relate to aperiodic neural activity. Pearson correlations with cluster-based correction revealed that lower aperiodic exponents in both EC and EO conditions were generally associated with higher ADHD traits. Negative clusters were observed for ASRS total scores (EO: p = .03; EC: p = .023; Fig. 2B, left), hyperactivity–impulsivity (EO: p = .002; EC: p > .05; Fig. 2B, middle), and inattentiveness (EO: p = .025; EC: p = .015; Fig. 2B, right). Together, these results indicate that individuals with higher ADHD traits may exhibit faster oscillatory sampling embedded within a noisier neural background.

**Figure 2.**
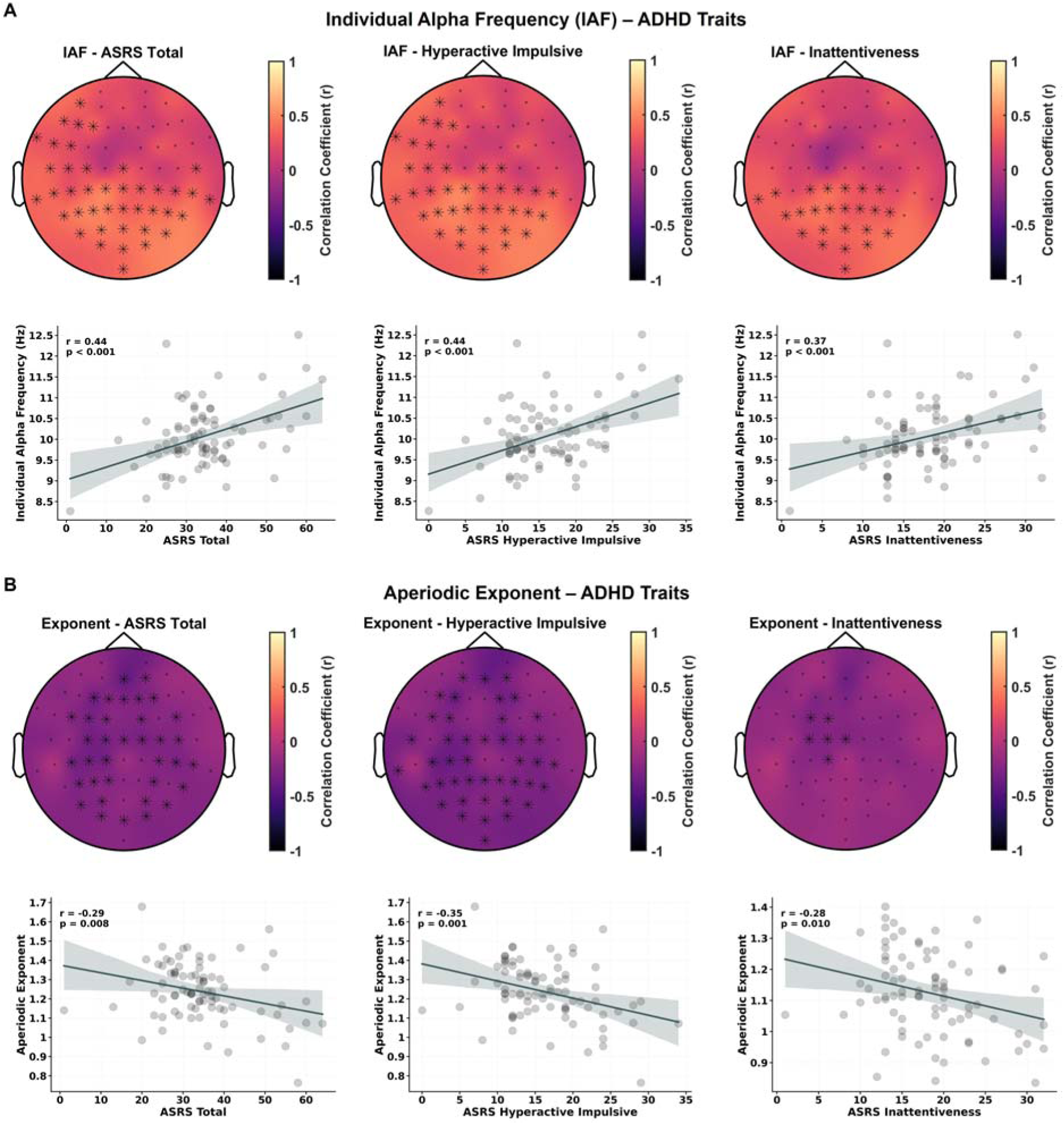
Alpha oscillatory speed and aperiodic neural activity predict ADHD traits. (A) Relationship between individual alpha frequency (IAF) and ADHD traits. Top row: scalp maps display electrode-wise Pearson correlations between resting-state IAF and ASRS total scores (left), hyperactivity–impulsivity (middle), and inattentiveness (right) in the eyes-open condition. Asterisks mark electrodes contributing to significant positive clusters identified by cluster-based permutation tests (10,000 permutations). Bottom row: scatterplots show individual participant data with regression lines and 95% confidence intervals. Faster IAF was associated with higher ADHD traits across all ASRS dimensions. (B) Relationship between aperiodic exponent and ADHD traits. Top row: scalp maps show electrode-wise correlations between the aperiodic exponent and ASRS total scores (left), hyperactivity–impulsivity (middle), and inattentiveness (right) in the eyes-open condition. Asterisks mark electrodes forming significant negative clusters. Bottom row: scatterplots depict individual datapoints, regression lines, and 95% confidence intervals. Lower aperiodic exponents, reflecting increased neural noise, were associated with higher ADHD traits. Together, these results indicate that individuals with higher ADHD trait levels exhibit faster alpha oscillatory speed and reduced aperiodic exponent values, consistent with faster neural sampling occurring within a noisier neural background.

### Neural predictors of atypical temporal perception in ADHD profiles

Our results showed that individual differences in ADHD traits were predicted by both neural measures and behavioural indices of temporal perception, including temporal thresholds (PSE) and their modulation by prior perceptual experience (ΔPSE). To determine whether neural dynamics accounted for the association between ADHD traits and temporal perception, we conducted mediation analyses to test whether these relationships were independent or mediated by neural measures. Path coefficients were estimated using maximum likelihood, and statistical significance was evaluated with 10,000 bias-corrected bootstrap resamples (see Methods).

We first tested whether IAF and aperiodic exponent mediated the association between ADHD traits and temporal thresholds PSE using parallel multiple mediation (Fig. 3A). The model showed excellent fit (χ² = 0.322, CFI = 1.00, TLI = 1.06, SRMR = 0.016) and explained 41.8% of the variance in temporal thresholds. ADHD traits were positively associated with faster IAF (a□ = 0.441, p < .001) and negatively associated with the aperiodic exponent (a□ = −0.290, p = .037). In turn, faster IAF predicted narrower temporal windows (b□ = −0.397, p < .001), whereas the path from the aperiodic exponent to temporal thresholds was not reliable (b□ = 0.174, p = .094). Accordingly, the indirect effect of ADHD traits on temporal thresholds via IAF was significant (a□b□ β indirect = − 0.175, BCa 95% CI [−0.353, −0.059]), whereas the indirect pathway via the aperiodic exponent was not (BCa 95% CI [−0.169, 0.001]). A residual direct effect of ADHD traits on temporal thresholds remained after accounting for both neural mediators (c′ = − 0.284, p = .009), yielding a significant total effect (−0.510, BCa 95% CI [−0.715, −0.245]). Additionally, to test whether joint oscillatory–aperiodic dynamics contributed beyond their main effects, we added an IAF × aperiodic exponent interaction to the outcome model. Model fit remained excellent (χ² = 0.816, CFI=1.00, TLI=1.10, SRMR=0.021), but neither the interaction term (b□ = 0.104, p = .193) nor the mediated interaction effect was significant (BCa 95% CI [−0.046, 0.002]). The indirect effect via IAF remained robust, indicating that IAF is the primary neural mediator of the ADHD–temporal threshold association.

**Figure 3.**
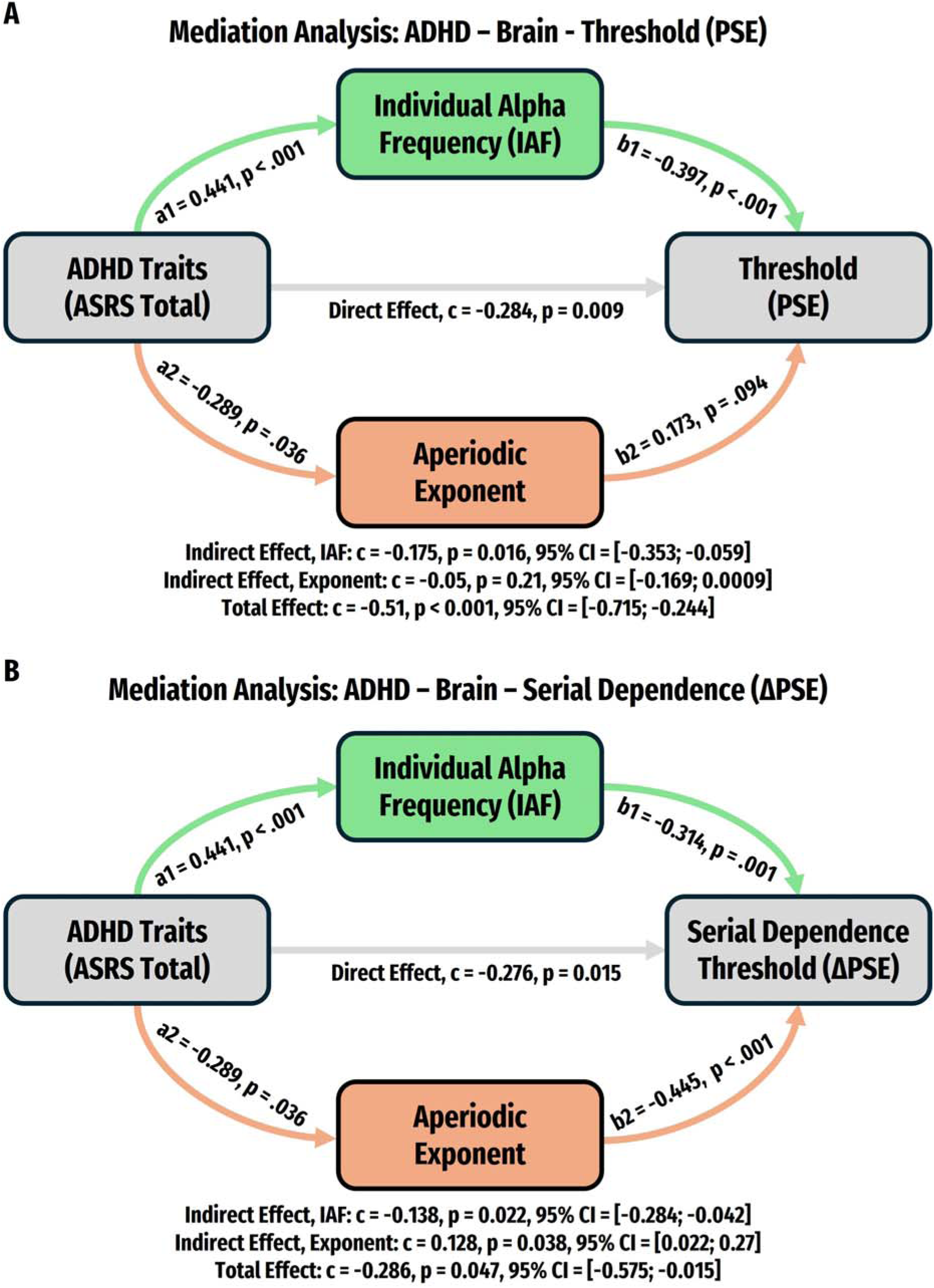
Neural predictors of temporal perception in ADHD-like profiles. **(A)** Multiple mediation model testing whether individual alpha frequency (IAF) and the EEG aperiodic exponent mediate the association between ADHD trait severity (ASRS total score) and temporal perception thresholds (point of subjective equality; PSE). ADHD traits were positively associated with faster IAF and negatively associated with the aperiodic exponent (a-paths). Faster IAF predicted narrower temporal windows (b□), whereas the aperiodic path to PSE did not reach significance (b□). The indirect effect via IAF was significant, whereas the indirect effect via the aperiodic exponent was not. A residual direct effect of ADHD traits on temporal thresholds remained after accounting for neural mediators. Path coefficients represent standardized estimates. **(B)** Parallel multiple mediation model testing whether IAF and the aperiodic exponent mediate the association between ADHD traits and serial dependence magnitude (ΔPSE), reflecting the influence of prior perceptual experience on current temporal judgments. ADHD traits were again associated with faster IAF and lower aperiodic exponent. Faster IAF predicted reduced serial dependence, whereas a lower aperiodic exponent predicted increased serial dependence, yielding two significant indirect effects. The direct ADHD–ΔPSE path remained significant, while the total effect was modest due to partial cancellation between the competing indirect pathways. No evidence was found for synergistic or moderated mediation, as inclusion of an IAF × aperiodic exponent interaction did not improve model fit or explained variance. Indirect effects were tested using 10,000 bias-corrected bootstrap resamples and were considered significant when the 95% confidence interval did not include zero.

We next examined the same mediation structure for assessing the effects of prior perceptual experience (serial dependence magnitude: ΔPSE; Fig. 3B). The parallel mediation model showed excellent fit (χ² = 0.322, CFI = 1.00, TLI = 1.08, SRMR = 0.017) and accounted for 33.8% of the variance in ΔPSE. ADHD traits were associated with faster IAF and a lower aperiodic exponent, replicating the neural paths observed for temporal thresholds. Faster IAF predicted reduced serial dependence (b□ = − 0.314, p = .002), whereas a lower aperiodic exponent predicted increased serial dependence (b□ = − 0.445, p < .001). Both indirect effects were significant but operated in opposite directions: a negative indirect effect via IAF (BCa 95% CI [−0.284, −0.042]) and a positive indirect effect via the aperiodic exponent (BCa 95% CI [0.022, 0.271]), indicating that ADHD-related differences in oscillatory timing and broadband neural excitability exert significant influences on perceptual history biases. The direct ADHD–behavior path remained significant (c′ = −0.277, p = .016), while the total effect was modest. As for temporal thresholds, adding an IAF × aperiodic exponent interaction did not improve model fit or explained variance and provided no evidence for synergistic mediation. Together, these findings indicate that oscillatory and aperiodic neural dynamics independently mediate ADHD-related variability in temporal perception, with converging effects on temporal thresholds but opposing influences on serial dependence.

## Discussion

Atypical temporal processing is thought to represent a core component of the neurocognitive architecture of ADHD, with individuals often perceiving time as passing unusually quickly^15^ and showing anomalies in integrating the continuous stream of sensory input. Such distortions in temporal experience are believed to contribute to hallmark features of the disorder, including difficulties with sustained attention and hyperactive–impulsive behaviour^14,15^. Although temporal processing has long been linked to the intrinsic speed of alpha-band oscillations^25,27,28,33^ and, more recently, to non-rhythmic aperiodic neural dynamics^27,28,29,65^, their role in ADHD remains unexplored. Here, we show that resting-state oscillatory and aperiodic neural activity vary systematically with ADHD trait levels in the general population and jointly predict the temporal perception profile associated with elevated ADHD features.

First, our findings revealed that individuals with higher ADHD traits exhibit a narrower window of temporal binding for visual information, tending to overestimate temporal asynchrony. This pattern aligns with previous evidence of atypical temporal integration across both uni- and multisensory domains in individuals with elevated ADHD features^18,19,22,23,24^. Increased perceived temporal misalignment across sensory events may contribute to the characteristic behavioural phenotype of distractibility and inattentiveness observed in this condition^22,23,24^. Consistent with prior work in neurotypical samples^23^, a smaller temporal integration threshold was similarly associated with both the inattentiveness and the hyperactivity–impulsivity ADHD dimension, indicating that atypical temporal integration represented a shared feature across symptom domains in our sample.

Importantly, we also found that individuals with higher ADHD traits showed reduced serial dependence in their temporal judgments, indicating diminished flexibility in adaptively shaping temporal processing based on recent perceptual experience. In the general population, temporal integration judgments are typically biased toward the previous perceptual outcome, reflecting an adaptive mechanism that maintains perceptual continuity across time^7,29,47,48^. The reduced perceptual history dependence observed in individuals with elevated ADHD traits may reflect excessive precision assigned to incoming sensory information, combined with increased attentional fluctuations and a faster decay of information over short time scales^48,49,50,51,52,53,54,55,56,57,58,59,60^. These factors that may weaken reliance on prior internal models often reported in ADHD profiles, ultimately leading to atypical temporal processing.

But how is this distinctive temporal processing profile supported at the neural level? First, our findings revealed that higher ADHD traits were associated with faster individual alpha frequency (IAF) in both eyes-open and eyes-closed resting-state conditions. More specifically, faster IAFs were observed over posterior–central scalp regions for individuals with elevated anomalies in inattentiveness and hyperactivity–impulsivity dimensions. Notably, faster IAF has been consistently linked to heightened arousal, increased vigilance, and faster information processing^66,67,68^, in line with the hyper-responsive and over-activated cortical states often described in ADHD. These results extend prior research that has predominantly examined differences in alpha-band power in ADHD^69,70,71^, suggesting that alpha oscillatory speed may represent a useful resting-state signature of ADHD-related profiles.

Additionally, our results indicated a generally lower aperiodic neural exponent in individuals with higher ADHD traits. This finding aligns with recent reports showing a markedly reduced aperiodic exponent in ADHD^39,40^, a pattern reflecting increased neural noise and altered cortical excitation–inhibition balance^72^, but also brain states characterised by vigilance and arousal^73,74^, potentially linked to dysregulation of dopaminergic and noradrenergic systems^41,42,43,44^. Notably, the scalp distribution of this atypical aperiodic neural dynamic revealed dissociations across ADHD domains. Lower aperiodic exponent values associated with hyperactive–impulsive features were spatially widespread, whereas associations with inattentive traits were more restricted to central scalp regions. This pattern may potentially suggest differential involvement of large-scale cortical processes across ADHD dimensions. In line with prior evidence, hyperactivity–impulsivity may be linked to more globally distributed alterations in cortical excitability, whereas inattentiveness may involve more spatially circumscribed neural differences related to attentional control processes^75,76,77^.

Critically, our results show that these neural signatures influenced visual temporal performance across different levels of ADHD trait expression. Mediation analyses revealed that individuals with elevated ADHD traits exhibited systematically faster IAF, which in turn predicted narrower temporal integration windows. These findings align with prior frameworks proposing that alpha activity provides a temporal scaffold for perception, with faster alpha rhythms supporting narrower temporal binding windows^25^. Within this view, our results suggest that ADHD-related alterations in perceptual timing may be partly instantiated through faster neural oscillatory sampling of visual input that constrain the temporal window over which sensory information is integrated.

Furthermore, our findings showed that oscillatory and aperiodic neural dynamics jointly determined how strongly recent perceptual experience shapes current temporal judgments across individual differences in ADHD. Overall, faster IAF and a lower aperiodic exponent were associated with reduced serial dependence, supporting the idea that faster alpha rhythms operating within a less noisy neural background may promote more accurate sampling of visual input and yield a more reliable representation of sensory events^27,28,29^. Critically, in individuals with high ADHD traits, faster IAF accounted for the reduced influence of prior perceptual outcomes, suggesting that faster oscillatory sampling may bias perception toward heightened moment-to-moment sensory processing at the expense of perceptual stability. In parallel, higher ADHD traits predicted a flatter aperiodic exponent, and increased neural noise in turn predicted stronger perceptual history dependence. This aperiodic pathway reflects an alternative mechanistic route through which ADHD-related increases in neural noise amplify reliance on prior perceptual experience, counteracting the history-reducing effect of faster alpha oscillations, suggesting that individual differences in perceptual history biases reflect the net outcome of competing neural processes.

Together, these findings argue against a single-mechanism account of temporal processing in ADHD traits and instead support a multidimensional neural model in which oscillatory alpha frequency constrains the temporal window of integration and updating speed, whereas aperiodic neural dynamics support perceptual continuity by regulating the persistence of internal representations rather than temporal integration per se.

## Methods

### Participants

A total of 83 individuals took part in the study (46 females; mean age = 23.4 years, SD = 2.58). All participants reported normal or corrected-to-normal eyesight. Individuals with a history of neurological conditions, epilepsy or photosensitivity were excluded based on self-report. The behavioural task data from the present work had already been included in a previous study (Marsicano et al., 2025). The target sample size was determined a priori based on a recent meta-analysis of 27 studies examining the association between individual alpha frequency and temporal resolution^25^ (see also^29^). All procedures adhered to the Declaration of Helsinki and were approved by the Institutional Review Board of the New York University Abu Dhabi Human Research Protection Program.

### ADHD Traits

ADHD traits were quantified using the Adult ADHD Self-Report Scale (ASRS^62^). The ASRS consists of 18 items assessing the frequency of DSM-IV-TR Criterion symptoms of adult ADHD, rated on a five-point scale from never (0) to very often (4). The questionnaire reliably captures two symptom dimensions — a nine-item inattention factor and a nine-item hyperactivity–impulsivity factor — supported by previous factor-analytic work^78^. Reported internal consistencies are adequate for both subscales (α = 0.75 for inattention; α = 0.77 for hyperactivity–impulsivity) as well as for the total scale (α = 0.82). For each participant, we computed total ASRS scores alongside inattentiveness and hyperactivity–impulsivity subscale scores, with higher values reflecting greater expression of ADHD profiles. The ASRS is widely used in both clinical and general-population research as a reliable screening measure^79^.

### Stimuli and Experimental Design

As described in our recent study^29^, we used a visual temporal integration paradigm designed to estimate individual integration performance while avoiding methodological limitations of conventional brief two-flash tasks^61^. Stimuli were generated in MATLAB (MathWorks) using Psychtoolbox and displayed on a gamma-corrected monitor operating at 144 Hz. Participants viewed the screen binocularly from 70 cm in a dimly lit testing room. Each trial consisted of two Gabor patches presented at the same central location and alternated over approximately 1 s. The Gabors had orientations of +45° and −45°, a spatial frequency of 1.94 cycles/degree (λ = 0.52°), a Gaussian envelope (σ = 0.62°), and full contrast. Each patch was rendered within a circular aperture and converted into a texture. Temporal perception was manipulated by varying the duration of each Gabor’s presentation (inter-stimulus interval, ISI), rather than by inserting temporal gaps between discrete flashes. The two orientations alternated in counterphase with ISIs ranging from 0 to 146 ms in 21-ms increments. Each ISI was presented 30 times, for a total of 240 randomly ordered trials. When ISI = 0, a precomputed fused stimulus with subtle spatial jitter was shown to prevent adaptation. The orientation that appeared first on each trial was randomized. This design created a continuous sequence in which short ISIs produced the percept of a single fused plaid (reflecting temporal integration), whereas longer ISIs led to the perception of two alternating gratings (reflecting temporal segregation). Because both orientations were spatially overlapping and presented without intervening gaps, the paradigm isolates genuine temporal binding rather than detection failures. At the start of the session, resting-state EEG was recorded for 3 min with eyes closed (EC) and 3 min with eyes open (EO). Participants then completed the ASRS questionnaire, followed by task instructions and a brief practice block. Each trial began with a fixation display (700–1300 ms), followed by the ∼1-s alternation sequence. After stimulus offset, participants provided a two-alternative forced-choice response indicating whether they perceived a single fused pattern (key “F”) or two alternating gratings (key “S”). Participants also provided a four-point confidence judgment^29^, however, these data are not included in the present report because they are beyond the aims of this study. A short break was offered halfway through the experiment.

### Data Analysis

#### Behavioral and Psychometric Function Analyses

Regarding the behavioral performance, for each participant, we quantified temporal integration performance by calculating the proportion of “separate” responses for every inter-stimulus interval (ISI)^29^. These data were then modeled with an individual psychometric curve, using a logistic function fit to the proportion of segregation reports across ISIs. Curve fitting was performed with a nonlinear least-squares procedure, yielding high-quality fits across observers (mean adjusted R² = 0.938; all participants > 0.85). The formula applied was as follows: *y = 1/(1 + exp(b × (t - x)),* where x denotes the ISI and y the predicted proportion of separate percepts. The function was constrained between 0 and 1, with two free parameters: the slope (b) and the temporal threshold (t), both restricted to positive values. From these fits, we extracted for each participant the 50% point of the function (threshold *t*) and the slope (*b*). The threshold corresponds to the Point of Subjective Equality (PSE), indexing the timing at which fused and separate percepts are equally likely. Lower PSE values indicate a stronger bias toward segregation (i.e., a narrower temporal integration window), whereas higher PSEs reflect increased temporal binding. The slope parameter captures the sharpness of the transition from fused to separate percepts, serving as an index of temporal precision: larger slopes correspond to more reliable, less noisy perceptual performance. To examine how perceptual history shaped current judgments, we implemented a serial-dependence analysis based on the percept reported on the immediately preceding trial. Trials were classified into two categories: those following a segregation percept (t–1 separate) and those following an integration percept (t–1 fused). Separate logistic functions were then fit for the two history conditions (mean adjusted R² = 0.918 for t–1 fused trials; 0.92 for t–1 separate trials). This yielded distinct PSE and slope estimates for each bin. Serial dependence strength was quantified as the PSE difference between PSE t–1 integration – PSE t–1 segregation bins (ΔPSE = PSE t–1 integration – PSE t–1 segregation), with positive values indicating an attractive bias toward the previous percept. In parallel, we computed an analogous measure for the slope parameter (Δslope) to assess whether perceptual history also influenced temporal precision.

### EEG analysis

Resting state EEG was recorded in two conditions, eyes closed (EC) and eyes open (EO), each lasting 3 minutes. Signals were acquired from 64 active scalp electrodes using BrainVision Recorder (Brain Products), sampled at 1000 Hz and referenced online to FCz. Electrode impedances were kept below 10 kΩ. Participants were instructed to relax, minimize movements, and either fixate a central point during the EO condition or keep their eyes closed during the EC condition. Preprocessing was performed in MATLAB. For each participant, electrodes containing noise were identified by visual inspection and replaced using spherical spline interpolation. Oculomotor activity was removed using independent component analysis (ICA). The continuous recordings were then segmented into 1 second epochs, and artifact free segments were submitted to spectral decomposition with Welch’s method to obtain power spectral density (PSD) estimates. Following our recent work^27,28,29^, aperiodic profiles were characterized using the FOOOF algorithm (Fitting Oscillations and One Over F^64^). This algorithm models the log power spectrum as the combination of an aperiodic component, described by an offset and an exponent, and a set of Gaussian peaks representing rhythmic activity. Spectra were fitted in the 1 to 70 Hz range to avoid edge related distortions. All fits used the default parameter settings (peak width limits 0.5 to 12 Hz, minimum peak height 0, no limit on the number of peaks, peak threshold 2), consistent with prior studies^64^. The aperiodic exponent derived from this fit served as an index of aperiodic neural activity, which has been linked to cortical excitation to inhibition balance and temporal perception^27,28,29,37,72^. Individual alpha frequency (IAF) was determined as the frequency corresponding to the maximum periodic peak within the 8 to 13 Hz range.

### Statistical Analysis

#### Link Between ADHD Traits and Temporal Perception

To examine the association between visual temporal perception and ADHD traits, we first computed Pearson correlations between ASRS scores and the psychometric parameters derived from the logistic fits (temporal threshold, PSE; and slope, b). These analyses were conducted separately for the ASRS total score, the inattention subscale, and the hyperactivity–impulsivity subscale. We then assessed whether ADHD traits modulated the influence of prior perceptual judgments by correlating ASRS scores with the serial-dependence indices derived from the behavioral task (ΔPSE and Δslope). These measures quantify the extent to which percepts on the preceding trial biased subsequent temporal judgments. In all analyses, ADHD trait scores were treated as continuous variables. Rather than categorizing individuals into “high” and “low” trait groups, we followed methodological recommendations emphasizing that the continuous treatment of dimensional measures preserves variance, maintains statistical power, and avoids the information loss and inflated Type II error rates associated with dichotomization^63,80,81,82^. For visualization purposes only, and not for inferential testing, we additionally generated illustrative plots (Fig. 1B–C) based on a median split of the ASRS total scores, yielding two groups: a low-trait subgroup (n = 48) and a high-trait subgroup (n = 35).

### Link Between ADHD Traits and Neural Activity

We examined the relationship between the neural measures of interest (IAF and aperiodic exponent), and ADHD traits. Analyses were performed separately for the eyes closed (EC) and eyes open (EO) resting state conditions. For each condition, we computed Pearson correlations between ADHD trait scores and the neural measures at every electrode, and statistical significance was assessed using nonparametric cluster-based permutation tests to control for multiple comparisons (10.000 permutations^83^). These analyses were carried out separately for the ASRS total score, the inattention subscale, and the hyperactivity and impulsivity subscale. In the present report we do not include results comparing EC and EO conditions, nor do we report analyses examining associations between the neural measures and the behavioral indices from the temporal perception task across the full sample irrespective of ADHD traits. These findings have already been reported in previous work (Marsicano et al., 2025) and are not central to the aims of the current study, which focuses on the relationship between ADHD traits and neural markers of visual temporal perception. All EEG analyses were performed in MATLAB using custom code together with the FieldTrip Toolbox^84^. Spectral parameterization was carried out with the FOOOF-mat toolbox^64^ running within a Python 3 environment.

### Mediation Analyses: ADHD Traits-Neural Activity-Temporal Perception

To examine whether neural dynamics accounted for the association between ADHD traits and temporal perception, we conducted mediation analyses using the *lavaan* package (version 0.6–19) in RStudio. All variables were standardized prior to analysis. Structural equation models were estimated with maximum likelihood and Full Information Maximum Likelihood to handle missing data. The significance of indirect effects was evaluated using 10.000 bias corrected bootstrap resamples, from which 95% confidence intervals were obtained. All reported coefficients are standardized. We first tested whether individual alpha frequency (IAF) and the aperiodic exponent mediated the relationship between ADHD traits and temporal thresholds (PSE). A parallel mediation model was specified in which ADHD traits predicted both neural measures, and PSE was predicted simultaneously by IAF, the aperiodic exponent, and ADHD traits. This model provided estimates of the indirect effect through IAF, the indirect effect through the aperiodic exponent, the direct effect of ADHD traits on PSE, and the total effect. We then evaluated an interaction augmented model to determine whether the combined EEG profile, defined as the product of standardized IAF and aperiodic exponent values, further accounted for variance in PSE. In this model, PSE was predicted by IAF, the aperiodic exponent, their interaction term, and ADHD traits. This approach allowed us to quantify a mediated interaction effect, reflecting whether ADHD traits influenced PSE through the joint contribution of oscillatory and aperiodic activity. Model comparison focused on the change in explained variance relative to the parallel mediation model. All models were tested on EEG values averaged over posterior electrode clusters showing significant correlations in the eyes-open (EO) condition. Effects were considered significant when the 95% bootstrap confidence interval of the indirect effect did not include zero. Model fit was evaluated using standard indices (CFI, TLI, RMSEA, SRMR), and explained variance (R²) was reported for each dependent variable. We performed a second set of mediation analyses replacing PSE with the magnitude of serial dependence, quantified as the difference in temporal thresholds between trials preceded by fused versus separate percepts (ΔPSE). The same model structure was applied, with ADHD traits as the predictor, IAF and the aperiodic exponent as mediators, and ΔPSE as the outcome. Parallel and interaction augmented models were estimated, allowing us to test whether oscillatory dynamics, aperiodic activity, or their combined profile accounted for individual differences in the influence of perceptual history on current temporal judgments. Estimation, inference, and model evaluation followed the same procedures described for the PSE models, and standardized indirect, direct, and total effects were reported.

## Acknowledgements

This work was supported by the NYUAD Center for Brain and Health, funded by Tamkeen under NYU Abu Dhabi Research Institute grant CG012.

## Author contributions

Conceptualization, G.M., M.D., D.M.; Methodology G.M., M.D., D.M.; Formal Analysis, G.M.; Software, G.M.; Visualization, G.M.; Funding Acquisition, D.M.; Writing - Original Draft Preparation, G.M. D.M.; Writing – Review & Editing, G.M., M.D., D.M.; Supervision, D.M.

