## Supplementary Materials for "Oscillatory and aperiodic neural dynamics shape temporal perception and weighting of perceptual priors in ADHD profiles"

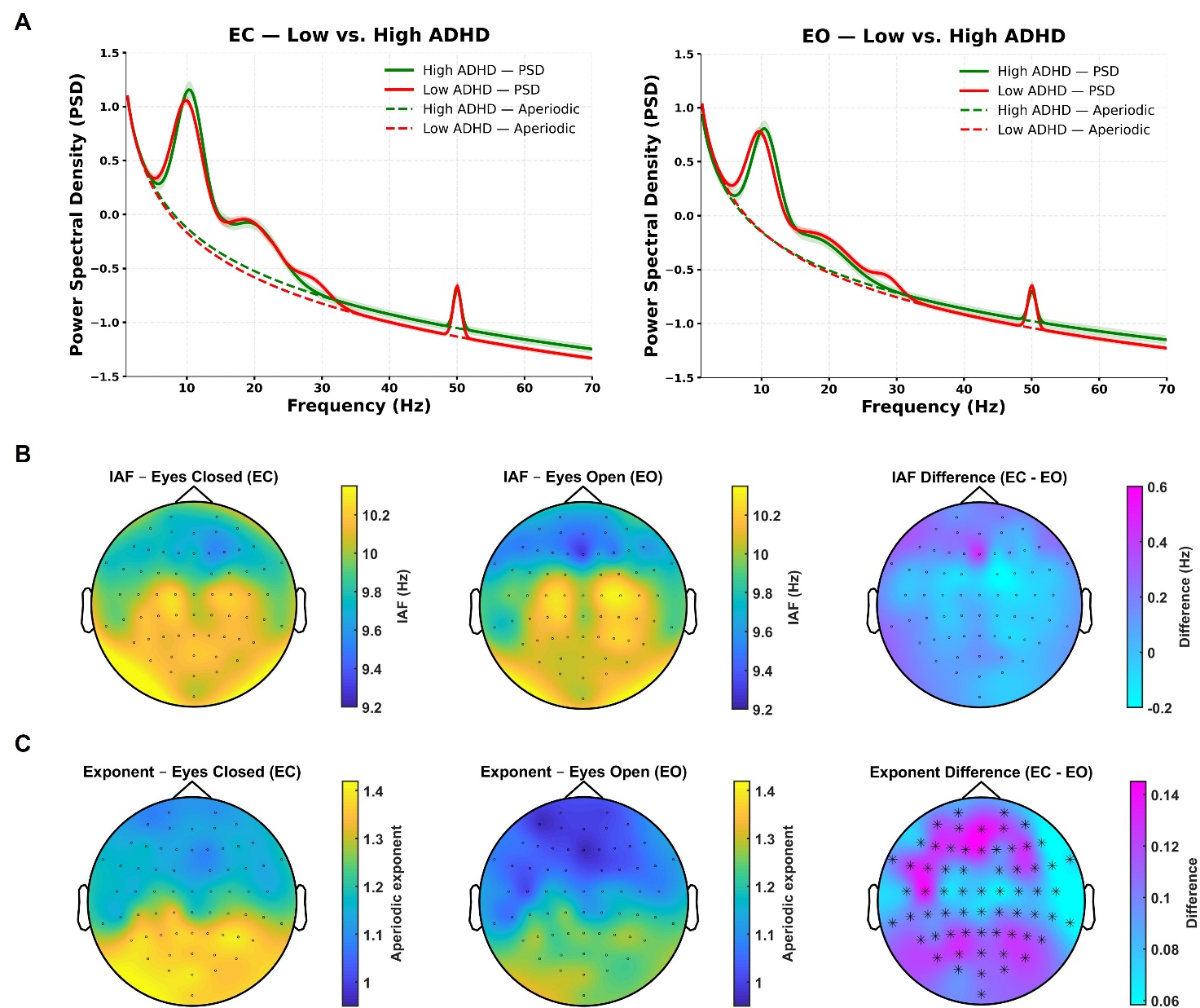

**Figure S1. Resting-state EEG decomposition and comparison of oscillatory and aperiodic neural activity across eyes-closed and eyes-open conditions.** (A) Group-averaged FOOOF spectral parameterization of resting-state EEG power spectra for individuals with low (red) and high (green) ADHD traits, defined using a median split of ADHD traits (ASRS total scores), during eyes-closed (EC, left) and eyes-open (EO, right) conditions. Power spectral density (PSD; solid lines) is averaged across posterior–central electrodes showing highest associations with ADHD trait scores and spectral measures (see results). The FOOOF algorithm decomposes the PSD into its aperiodic (1/f) component (dashed lines) and periodic oscillatory peaks. Shaded regions indicate ± SEM across participants. (B) Topographical distributions of IAF during EC (left) and EO (middle) resting-state conditions, and their scalp-level difference map (EC–EO, right). No significant differences in IAF were observed between conditions (p > 0.05, cluster-corrected). (C) Topographical distributions of the aperiodic exponent during EC (left) and EO (middle), and the corresponding difference map (EC–EO, right). Aperiodic exponents were significantly higher during EC relative to EO (p < 0.001, two-tailed, cluster-based correction; black asterisks indicate significant electrodes), consistent with stronger aperiodic activity under eyes-closed rest.
